# The Role of Parvalbumin-positive Interneurons in Auditory Steady-State Response Deficits in Schizophrenia

**DOI:** 10.1101/662510

**Authors:** Christoph Metzner, Bartosz Zurowski, Volker Steuber

## Abstract

Despite an increasing body of evidence demonstrating subcellular alterations in parvalbumin-positive (PV^+^) interneurons in schizophrenia, their functional consequences remain elusive. Since PV^+^ interneurons are involved in the generation of fast cortical rhythms, these changes have been hypothesized to contribute to well-established alterations of beta and gamma range oscillations in patients suffering from schizophrenia. However, the precise role of these alterations and the role of different subtypes of PV^+^ interneurons is still unclear. Here we used a computational model of auditory steady-state response (ASSR) deficits in schizophrenia. We investigated the differential effects of decelerated synaptic dynamics, caused by subcellular alterations at two subtypes of PV^+^ interneurons: basket cells and chandelier cells. Our simulations suggest that subcellular alterations at basket cell synapses rather than chandelier cell synapses are the main contributor to these deficits Particularly, basket cells might serve as target for innovative therapeutic interventions aiming at reversing the oscillatory deficits.

## Introduction

Since synchronized neuronal activity is thought to underlie efficient communication in the brain [8, 9, 31] and to be crucially involved in working memory processes [13], alterations of this synchrony, as found in electroencephalogram (EEG) or magnetencephalogram (MEG) studies of patients suffering from schizophrenia (SCZ) [33, 6], might contribute to the symptoms in patients and impairments characterizing schizophrenia. One very robust finding in EEG/MEG studies of schizophrenic patients is a deficit in the gamma band auditory steady state response (ASSR) [19, 41, 18, 36, 39, 22, 4, 34, 32, 23, 37].

Fast-spiking (FS), PV^+^, *γ*-amino-butyric acidergic (GABAergic) interneurons seem to be a major contributor to gamma oscillations [11, 1]. However, this class of inhibitory interneurons can be further divided into at least two subgroups: basket cells (BCs) and chandelier cells (ChCs) [15, 38]. The main difference between basket and chandelier cells is their axonal target. While basket cells mainly target the perisomatic region of excitatory pyramidal cells and other inhibitory interneurons, Chandelier cells exclusively target the axon hillock of pyramidal cells [15]. Furthermore, although their electrophysiological properties are similar subtle differences, such as different afterhyperpolarization amplitudes and firing frequencies, between the two subclasses exist [30]. Lastly, while BC synapses display fast and strong inhibition, ChC synapses have been shown to have a depolarizing effect *in vitro* [35, 44, 45, 46]. However, whether this depolarization also exists *in vivo* and whether it has an excitatory or rather a predominantly shunting effect remains elusive. Moreover, it is not clear so far whether these differences also imply a different involvement in the generation of gamma oscillations.

Several alterations of PV^+^ neurons have been found in schizophrenia. While the density of PV^+^ neurons does not seem to be altered in subjects with schizophrenia [43, 2], the levels of mRNA for the calcium-binding protein PV are markedly reduced [12]. Furthermore, 50% of PV^+^ neurons in dorsolateral prefrontal cortex lack detectable levels of the 67 kDa isoform of glutamate decarboxylase (GAD_67_), an enzyme that catalyzes the decarboxylation of glutamate to GABA [12]. It has been suggested that the reduced expression of GAD67 mRNA, the gene encoding the GAD_67_ enzyme, likely implies a reduction in GABA synthesis in cortical GABA neurons, which in turn would lead to smaller amplitude IPSCs at the postsynaptic site [10]. Furthermore, a reduction in the plasma membrane GABA transporter GAT1, which is a major contributor to the specificity of synapses by preventing spillover to neighbouring synapses [29, 16], has been found in PV^+^ interneurons in schizophrenic patients [43]. A reduction in GAT1 leads to a prolongation of IPSC durations [29, 16]. Interestingly, in patients with schizophrenia, immunoreactivity for the GABA-A *α*2 subunit is increased [42] in the axon initial segment of pyramidal cells, the exclusive target of chandelier cell cartridges. GABA receptors with an *α*2 subunit, compared to receptors with an *α*1 subunit, also show an increase in IPSC duration in tranformed human embryonic cells [20]. In summary, the reduced expression of GAD67, the redcution of GAT1 and increased α2 subunit expression in SCZ patients have two main effects: a reduction of inhibitory strength of PV^+^ neurons (both BCs and ChCs) and an increased IPSC duration at PV^+^ synapses (also for both BCs and ChCs, however, considerably longer for ChCs). These cellular differences in PV^+^ neurons are likely to have an influence on oscillatory activity. Since gamma oscillation generation and maintenance crucially depends on fast-spiking PV^+^ interneurons, changes in peak amplitude and duration of IPSCs might particularly affect oscillations in this frequency band.

Earlier modelling efforts by Vierling-Claassen et al. [41, 40], proposed that increased IPSC decay times at chandelier cell synapses might be sufficient to explain the gamma and beta range ASSR deficits in schizophrenia patients. However, while demonstrating that, in principle, increased IPSC decay times at chandelier cell synapses could result in experimentally observed ASSR deficits for patients with SCZ, Vierling-Claassen et al. did not consider the relatively small number of chandelier cells compared to basket cells [27]. Furthermore, they did not take into account potential increased IPSC decay times at basket cell synapses. Therefore, we extended their computational model to investigate the differential role of prolonged IPSC decay time at basket and chandelier cell synapses, at realistic ratios of chandelier to basket cells, in gamma and beta range ASSRs. We found that, while prolonged IPSC decay times at chandelier cells could explain experimental results when chandelier cells form 50% of inhibitory neurons in the network, they failed to do so at lower, more realistic percentages such as 10% or even 5%. In contrast, for physiologically plausible chandelier cell percentages of 10% or 5%, prolonged IPSC decay times at basket cell synapses were able to reproduce experimental findings. Furthermore, increased IPSC decay times at synapses of both interneuron subtypes resulted in qualitatively very similar ASSRs as for changes at basket cell synapses only, suggesting that chandelier cell alterations might not be crucially involved in gamma and beta range ASSR deficits in schizophrenia.

## Methods

### The Model

The model proposed here is based on a recent reimplementation [28] of the simple model presented by Vierling-Claassen et al. [41]. However, we extended the model to consist of two distinct inhibitory populations (instead of only one), one representing basket cells and one representing chandelier cells (see Figure 1).

**Figure 1:**
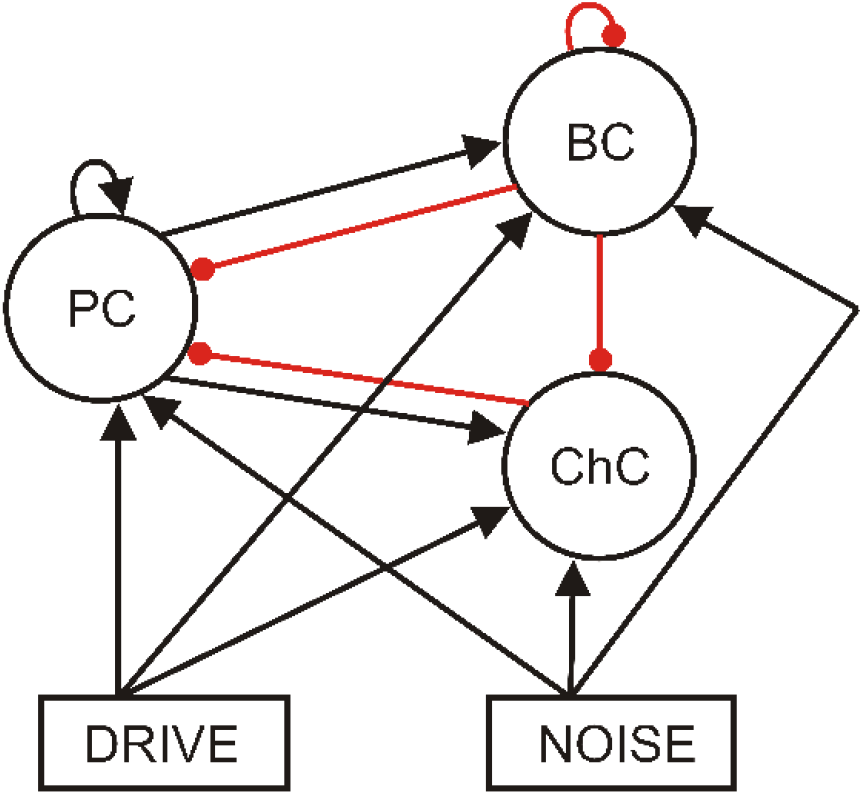
Schematic representation of the network structure. All three populations receive drive and noise input. The excitatory population makes connections to itself and to both inhibitory populations. Similarly, the basket cell population inhibits itself and both other populations. The chandelier cell population, however, exclusively targets the excitatory cells. (Black arrow: excitatory connection. Red arrow: inhibitory connection.)

#### Single Cell Model

Individual cells are modeled as theta neurons (for a detailed discussion of this neuron model see e.g. [3]).

The *k*th neuron in a network is described by a single variable *θ_k_*, which can be regarded as the neuron state, subject to the following dynamics

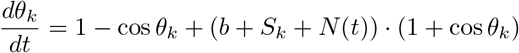

where *b* is an externally applied current, *S* is the total synaptic input to the cell and *N*(*t*) is a time-varying noise input. Total synaptic input to a cell in a network is calculated as

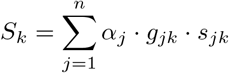

where *n* is the number of presynaptic neurons, *α_j_* controls excitation and inhibition, i.e. is +1 for excitatory synapses and −1 for inhibitory ones, *g_jk_* is the synaptic strength from cell *j* to cell *k* and *s_jk_* is the synaptic gating variable from cell *j* to cell *k*. Synaptic gating variables are subject to the following dynamics

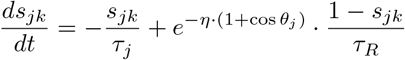

where *τ_j_* is the synaptic decay time, *τ_R_* the synaptic rise time and *η* is a scaling parameter. The network receives excitatory drive input at click train frequency from a single pacemaker cell. Additionally, Poissonian noise input is also given to all cells in the network. A noise spike at time *t_n_* elicits the following excitatory postsynaptic potential (‘EPSP’)

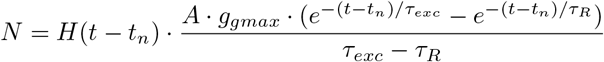

where *A* · *g_gmax_* is the strength of the noise, *τ_exc_* is the synaptic decay time, *τ_R_* the synaptic rise time, and *H* the Heaviside function.

#### Network

We combined 80 excitatory, pyramidal cells together with 40 inhibitory cells of the two different inhibitory subtypes (basket and chandelier cells) into a network model, following the earlier work of [41]. The percentages of the two different subtypes varied.

The connectivity of the network is summarized in Figure 1. Pyramidal cells make connections to other pyramidal cells, as well as to both types of inhibitory interneurons. Basket cells inhibit pyramidal cells, themselves and chandelier cells. Chandelier cells only inhibit pyramidal cells (see e.g. [38]). The connectivity between any two populations is all-to-all. All populations also have two sources of input, the oscillatory drive input and a background noise input. The drive input periodically sends spikes with a given frequency to all three populations. In order to generate the spikes a drive cell is implemented (also a theta neuron), which receives an applied current so that its spike frequency matches the driving frequency. This drive cell is then connected to all cells in the network. The background noise input sends noise spikes at times drawn from a Poisson distribution.

#### Entrainment Measures

In order to evaluate the oscillatory entrainment we record the spiking of the three populations as well as a simulated ‘EEG/MEG’ signal, calculated as the sum of all excitatory synaptic variables over all pyramidal cells [41]. As the main measure for entrainment we perform an FFT on the ‘EEG/MEG’ signal and extract the power in certain frequency bands. Since we are mainly interested in entrainment in the gamma and beta band, we specifically look at five measures (see experimental studies [19, 41]): 1) 40Hz power at 40Hz drive 2) 20Hz power at 40 Hz drive 3) 20 Hz power at 20 Hz drive 4) 40 Hz power at 20 Hz drive 5) 30 Hz power at 30 Hz drive (which will hereafter be referred to as the **40**/**40 measure**, the **20**/**40 measure**, the **20**/**20 measure**, the **40**/**20 measure** and the **30**/**30 measure**, respectively). As can be seen in Table 2, a decrease in the 40/40 measure is a robust finding in all three considered studies. Furthermore, the 30/30 measure is not affected in all three studies. Beyond the 40/40 measure, only the study by Vierling-Claassen et al. [41] found significant differences for other measures, namely, a decrease in the 40/20 measure, underpinning the deficit in gamma generation, and an increase in the 20/20 and the 20/40 measures, suggesting a transfer of power from the gamma to the beta band in patients with schizophrenia. It is important to note here, that both the Kwon et al. [19] and the Krishnan et al. [18] studies, used EEG whereas the Vierling-Claassen et al. [41] study used MEG and that it has been argued that MEG might have a higher sensitivity, which could explain the absence of findings other than for the 40/40 measure in the first two studies. Overall, given the experimental evidence discussed above, we think that the five measures chosen provide a good coverage of the frequency bands that are important to understand gamma and beta entrainment deficits in schizophrenia.

**Table 1:** Model parameters

| Parameter | Definition | Value |
| --- | --- | --- |
| $n_E$ | Exc. population size | 80 |
| $n_I$ | Inh. population size (BCs and ChCs together) | 40 |
|  | *Note that percentages of the two different types varies* |  |
| $\tau_R$ | Synaptic rise time | 0.1 |
| $\tau_{exc}$ | Excitatory decay time | 2.0 |
| $\tau_{bc}$ | BC decay time | 8.0 |
| $\tau_{chc}$ | ChC decay time | 8.0 |
| $g_{ee}$ | E-E synaptic strength | 0.00375 |
| $g_{ebc}$ | E-BC synaptic strength | 0.00625 |
| $g_{echc}$ | E-ChC synaptic strength | 0.00625 |
| $g_{bce}$ | BC-E synaptic strength | 0.00375 |
| $g_{chce}$ | ChC-E synaptic strength | 0.00375 |
| $g_{bcbc}$ | BC-BC synaptic strength | 0.005 |
| $g_{de}$ | Synaptic strength of drive to E cells | 0.3 |
| $g_{dbc}$ | Synaptic strength of drive to BC cells | 0.08 |
| $g_{dchc}$ | Synaptic strength of drive to ChC cells | 0.08 |
| $b$ | Applied current (regardless of cell type) | -0.1 |
| $Ag_{max}$ | Scaling factor for noise EPSCs | 0.6 |

**Table 2:** Summary of ASSR deficits in schizophrenic patients in the three studies considered here (↓: significantly lower in patients, ↑: significantly higher in patients, –: no significant difference between controls and patients).

| Exp. study | Measure |  |  |  |  |
| --- | --- | --- | --- | --- | --- |
|  | 40/40 | 30/30 | 20/20 | 40/20 | 20/40 |
| Kwon<br>et al.[19] | ↓ | - | - | - | - |
| Vierling-Claassen<br>et al. [41] | ↓ | - | ↑ | ↓ | ↑ |
| Krishnan<br>et al. [18] | ↓ | - | - | - | - |

#### Implementation

The model was implemented using Python 2.7.9 and numpy 1.9.3. Analysis and visualization of the model output was also done in Python using the numpy and matplotlib packages (matplotlib 1.4.3).

All differential equations were solved using a simple forward Euler scheme. A single simulation simulated a 500 ms trial and the time step was chosen such that this results in 2^13^ = 8192 data points. However, the main results were unaffected by using a smaller time step.

All code (for simulations, analysis and visualization) will be made publicly available in the following github repository: https://github.com/ChristophMetzner/Chandelier-Basket-Model.

### Exploration of Circuit Abnormalities

We explored the effects of natural differences between BCs and ChCs together with their differential schizophrenia-associated abnormalities on oscillatory entrainment in the gamma and beta frequency range. Specifically, we investigated the ratio of BCs versus ChCs and prolonged IPSC decay times of inhibitory synapses of both interneuron subtypes (because of a reduced expression of GAT1 for both BCs and ChCs and an increased expression of *α*_2_ subunits for ChCs). Table 3 details how these changes were modelled.

**Table 3:** Modelling of interneuron differences and circuit abnormalities

| Experimental finding | Parameter change in the model |
| --- | --- |
| BC/ChC ratio | $[n_{BC}, n_{ChC}]$: [20,20], [30,10], [36,4], [38,2] |
| Increased IPSC decay time at BC synapses | $\tau_{BC}$: 20 ms, 24 ms, 28 ms |
| Increased IPSC decay time at ChC synapses | $\tau_{ChC}$: 28 ms, 35 ms, 40 ms |

Because of the Poissonian background noise input, simulation results varied from trial to trial. To capture the robustness of findings we always performed 20 simulation trials, each with a different realisation of the underlying Poissonian noise process. When analysing the effect of circuit abnormalities, we averaged the 20 trials for each condition in time to get an average simulated MEG signal and calculated a power spectral density of this signal.

## Results

In order to explore the roles chandelier and basket cells play in the emergence of oscillatory entrainment deficits in schizophrenic patients, we simulated the effects of cellular and molecular abnormalities of both cell types found in schizophrenia. We started with an investigation of abnormalities of chandelier cells in isolation, followed by abnormalities of basket cells in isolation and concluded with an exploration of interactions of abnormalities in both cell types.

We always simulated the effect of abnormalities for four different conditions with a BC/ChC ratio of 50%, 25%, 10% and 5%, respectively. As results at high ChC percentages (50 and 25%) were similar and results at low ChC percentages (10 and 5%) were also similar, we only present results for 50% and 10%.

### Alterations of Chandelier Cells

As explained earlier, the plasma membrane GABA transporter GAT1 is reduced in chandelier cell synapses onto the axon initial segment of cortical pyramidal neurons, resulting in a prolongation of IPSC decay times. Additionally, the *α* subunit composition of the GABA_A_ receptors at the axon initial segment is skewed towards the *α*_2_ subunit in comparison to receptors at the dendrites, which leads to an even more prominent prolongation of decay times. This increased time course of inhibition at chandelier cell synapses has been proposed to account for the characteristic oscillatory entrainment changes in the gamma and beta range by Vierling-Claassen et al. [41, 40]. However, in their studies two important aspects of the local microcircuitry have not been accounted for: 1) the relatively small number of chandelier cells (compared to basket cells) [27] and 2) the specific connectivity scheme of chandelier cells, i.e. the fact that chandelier cells do not form connections with themselves [7]. We explored whether prolonged IPSC decay times at chandelier cell synapses (*τ_ChC_*=28 ms, as in [41]) could still explain schizophrenic entrainment deficits when these two aspects were included in the model.

At a ChC percentage of 50, the 40/40 measure was drastically reduced in the ‘schizophrenia-like’ network, reflecting a disturbance in the generation of gamma oscillations, which is in line with experimental evidence (see Table 2). However, the other entrainment measures did not match with experimental findings. The 30/30 measure was also strongly reduced, meaning that the disturbance in the generation of oscillations was not restricted to the gamma range at around 40 Hz. Furthermore, the 20/20 measure did not differ strongly between the ‘schizophrenia-like’ and the control network, while a difference was found in experimental studies (see [37]). Lastly, the emergence of a large 20/40 measure, which has been observed in the experimental study of Vierling-Claassen et al. as well as in their modelling work [41], could not be seen. As can be seen in Figure 2, the 20/40 measure, although higher than in the control network, remained small (note the scale in the 20/40 plot of Figure 2 (b)).

**Figure 2:**
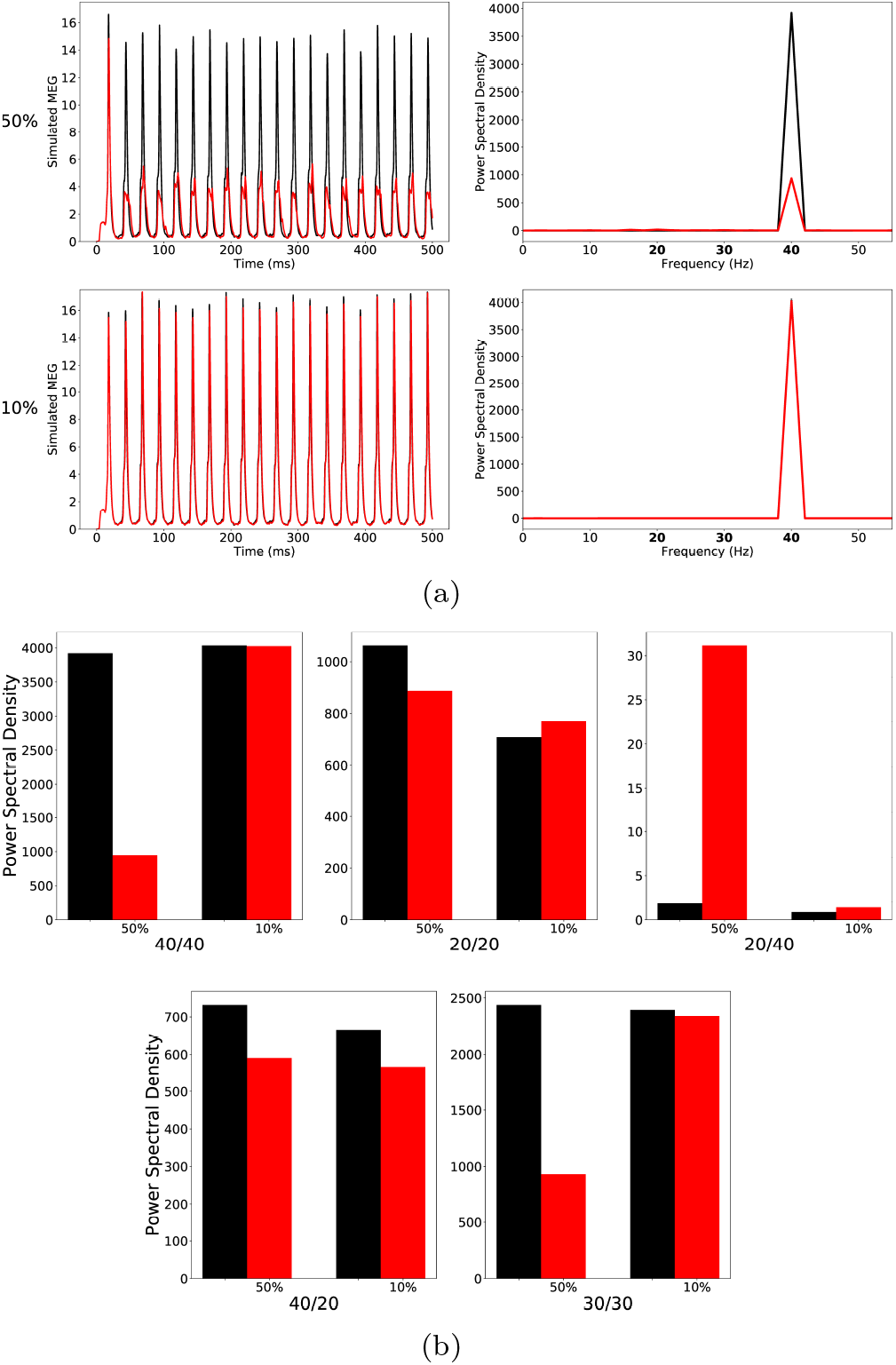
(a) left row: Raw simulated MEG signals for high (50%) and low (10%) percentage of chandelier cells among total inhibitory cells for a drive frequency of 40 Hz and the two network conditions (‘control’: black traces; ‘schizophrenia-like’: red traces). Right row: Power spectral density of the MEG signals of the left row. (b) Comparison of the 5 entrainment measures between control (black) and schizophrenia-like (red) for high and low ChC percentages. Note that ‘schizophrenia-like’ here means that IPSC decay times at **chandelier** cell synapses were increased from 8 ms to 28 ms. All traces here depict the mean signal averaged over 20 trials with different realisations of the Poisson process underlying the generation of noise. Importantly, averaging was performed in the time domain prior to a transformation into the frequency domain via Fourier transform, which is also the standard procedure in experiments.

At lower, more realistic, values of ChC percentage hardly any difference of the MEG signals and the five entrainment measures between control and ‘schizophrenia-like’ networks could be seen (see Figure 2). Note that this did not change with further increases of the IPSC decay times (we tested *τ_chc_* =35 ms and *τ_chc_* =40ms; data not shown). This suggests that prolonged IPSC decay times at ChC synapses are not sufficient to induce schizophrenia-like alterations in oscillatory entrainment.

### Alterations of Basket Cells

The plasma membrane GABA transporter GAT1 is also reduced in basket cell synapses onto dendrites in the perisomatic region of cortical pyramidal neurons, again resulting in a prolongation of IPSC decay times at these synapses [43]. However, opposed to chandelier cells, GABA_A_ receptors at basket cell synapses predominantly contain the α_1_ subunit [42]. This means that the IPSC decay time increases at BC-Pyr synapses probably also exist in schizophrenic patients but the increase is likely to be smaller than that at the ChC-Pyr synapses we examined before. Therefore, we initially chose an increased IPSC decay time of *τ_BC_* = 20 ms.

At high ChC percentages, this alterations had hardly any effect on the oscillatory behaviour of the network in response to any of the three driving frequencies tested (see Figure 3 (a) top row and Figure 3 (b)).

**Figure 3:**
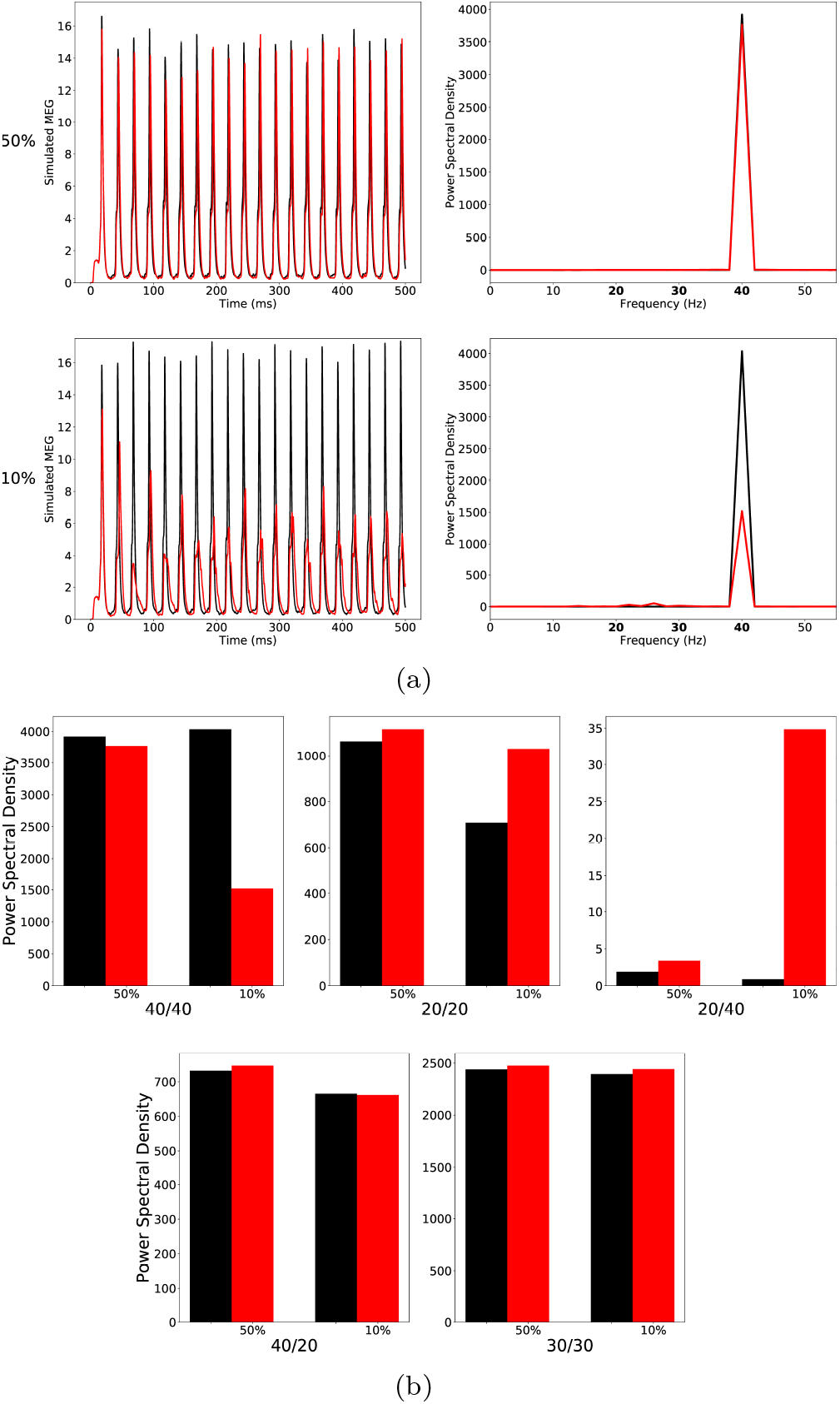
(a) left row: Raw simulated MEG signals for high (50%) and low (10%) percentage of chandelier cells among total inhibitory cells for a drive frequency of 40 Hz and the two network conditions (‘control’: black traces; ‘schizophrenia-like’: red traces). Right row: Power spectral density of the MEG signals of the left row. (b) Comparison of the 5 entrainment measures between control (black) and schizophrenia-like (red) for high and low ChC percentages. Note that ‘schizophrenia-like’ here means that IPSC decay times at **basket** cell synapses were increased from 8 ms to 20 ms. All traces here depict the mean signal averaged over 20 trials with different realisations of the Poisson process underlying the generation of noise. Importantly, averaging was performed in the time domain prior to a transformation into the frequency domain via Fourier transform, which is also the standard procedure in experiments.

At low, more realistic ChC percentages this alteration led to changes of oscillatory entrainment more in line with experimental evidence (see Figure 3). Specifically, the gamma range entrainment was signifcantly disturbed, reflected by the strong decrease of the 40/40 measure. Furthermore, a characteristic increase of the 20/20 measure was also clearly visible. The changes were restricted to the gamma and beta band leaving the respective border intact, i.e. the 30/30 measure was not significantly altered. However, the emergence of a strong 20/40 measure, seen by [41], both in experiment and model, was not visible. Figure 3 (b) (middle panel) does show an increase of the 20/40 measure compared to control, however it remains small.

Therefore, we increased the IPSC decay time at BC-Pyr synapses further to *τ_BC_* = 28ms. For high ChC percentages, the further increase of *τ_BC_* led to a strong decrease of the 40/40 measure but only had minor effects on all other entrainment measures (lower row of (a) and the third panel of (b) of Figure 4).

**Figure 4:**
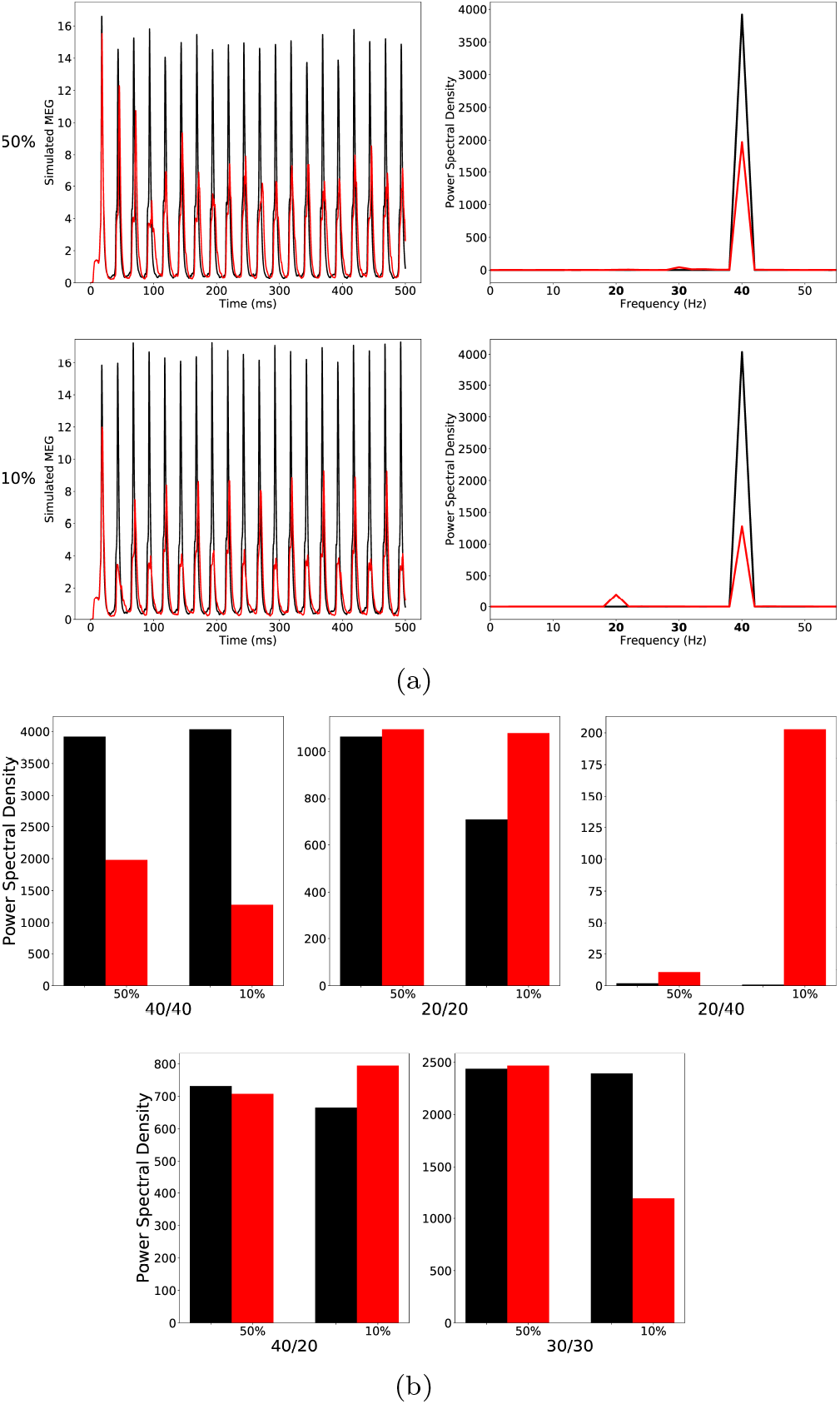
As Figure 3, but the IPSC decay times at **basket** cell synapses were increased to 28 ms to model ‘schizophrenia-like’ networks. (a) left row: Raw simulated MEG signals for high (50%) and low (10%) percentage of chandelier cells among total inhibitory cells for a drive frequency of 40 Hz and the two network conditions (‘control’: black traces; ‘schizophrenia-like’: red traces). Right row: Power spectral density of the MEG signals of the left row. (b) Comparison of the 5 entrainment measures between control (black) and schizophrenia-like (red) for high and low ChC percentages. All traces here depict the mean signal averaged over 20 trials with different realisations of the Poisson process underlying the generation of noise. Importantly, averaging was performed in the time domain prior to a transformation into the frequency domain via Fourier transform, which is also the standard procedure in experiments.

As shown in Figure 4 (lower row of (a) and the third panel of (b)), in the case of low ChC percentages, this led to a prominent increase of the 20/40 measure, similar to what was found in experiments [41]. Figure 5 (a) shows the evolution of this 20 Hz peak for 40 Hz drive over three different values of *τ_BC_* = 20 ms (black), 24 ms (red) and 28m s (green). While the Pyr cells are still able to recover from inhibition within every gamma cycle for *τ_BC_* = 20 ms, for larger values of *τ_BC_*, more and more Pyr cells are suppressed during every other gamma cycle, giving rise to a prominent 20 Hz peak. Interestingly, however, the longer IPSC decay time of *τ_BC_* = 28 ms did not only effect the 20/40 measure but also led to a strong decrease of the 30/30 measure, which then became markedly different from the 30/30 measure of the control network (see Figures 4 and 5 (b)). This is in disagreement with experimental findings.

**Figure 5:**
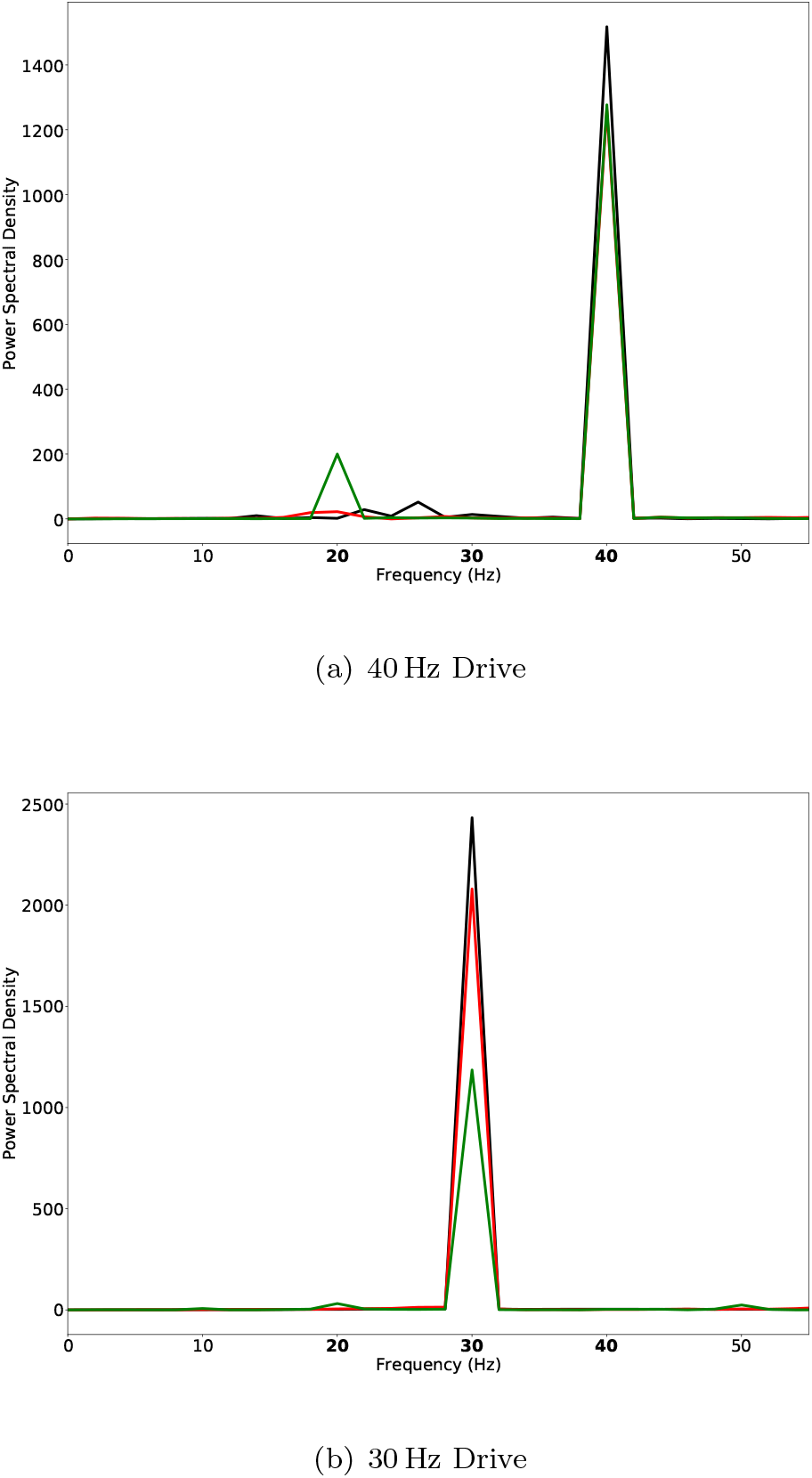
Power spectral density of simulated MEG signals for a ChC percentage of 10% for three different ‘schizophrenia-like’ networks with IPSC decay times at basket cells increased to 20 ms (black), 24 ms (red), and 28 ms (green), respectively. The network receives drive with a frequency of (a) 40 Hz and (b) 30 Hz.

### Alterations of Chandelier and Basket Cells

The results presented in the previous sections suggest that ChCs (at realistic numbers) are unable to produce ‘schizophrenia-like’ gamma and beta entrainment deficits in isolation and that BCs are the major contributor to these. However, alterations at ChCs might nevertheless modulate the deficits produced by BC alterations. Therefore, we next looked at the interactions of simultaneous alterations at BCs and ChCs.

First, we simulated the combined effect of a moderate increase of IPSC decay times at both ChC-Pyr and BC-Pyr synapses, with a stronger increase at ChC-Pyr synapses (i.e. *τ_ChC_* = 28ms and *τ_BC_* = 20ms).

Here, for high ChC percentages, we found a strong decrease of the 40/40 and the 40/20 measures (Figure 6), in line with experimental evidence of a gamma deficit in schizophrenia [19, 41, 37]. Furthermore, the 20/40 measure was increased, although still low. However, both the 20/20 and the 30/30 measure were also strongly reduced in this condition, in disagreement with experimental findings [19, 41]. For biologically realistic low ChC percentages we also found a strong reduction of the 40/40 measure (Figure 6 (a)), and a small reduction of the 40/20 measure (Figure 6 (b)). Interestingly, in this case the 20/20 measure was increased and, although still low, the 20/40 measure was also increased, while the 30/30 measure was only marginally different. Overall, the changes in spectral composition were consistent with experimental findings suggesting a general gamma deficit and beta increase in schizophrenia [41], similar to the case of *τ_BC_* = 20 ms.

**Figure 6:**
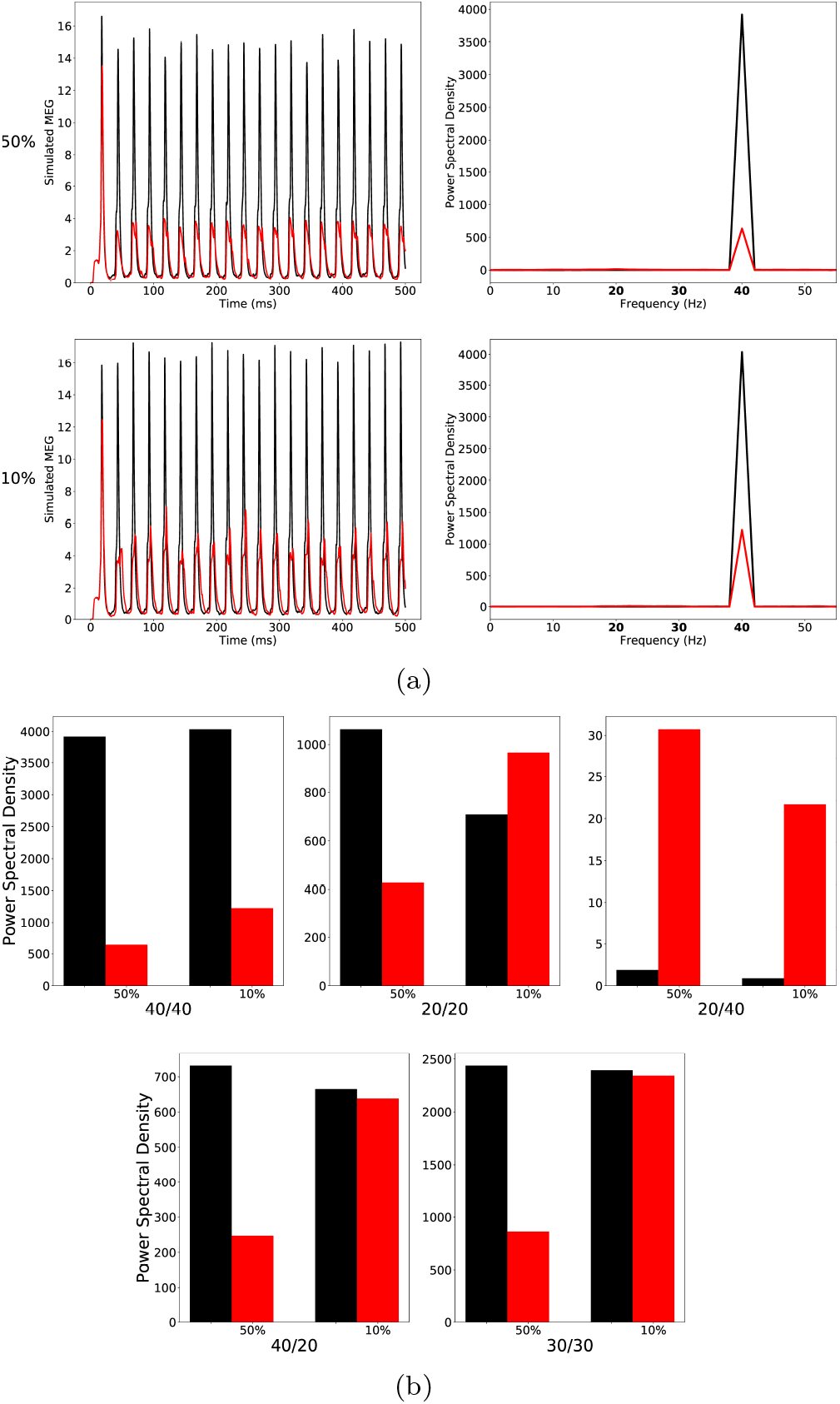
(a) left row: Raw simulated MEG signals for high (50%) and low (10%) percentage of chandelier cells among total inhibitory cells for a drive frequency of 40 Hz and the two network conditions (‘control’: black traces; ‘schizophrenialike’: red traces). Right row: Power spectral density of the MEG signals of the left row. (b) Comparison of the 5 entrainment measures between control (black) and schizophrenia-like (red) for high and low ChC percentages. Note that ‘schizophrenia-like’ here means that IPSC decay times at **chandelier** cell synapses were increased from 8 ms to 28 ms and at **basket** cell synapses were increased from 8 ms to 20 ms. All traces here depict the mean signal averaged over 20 trials with different realisations of the Poisson process underlying the generation of noise. Importantly, averaging was performed in the time domain prior to a transformation into the frequency domain via Fourier transform, which is also the standard procedure in experiments.

Lastly, we increased the IPSC times at BC-Pyr synapses further to 28 ms and, simultaneously, also increased IPSC times at ChC-Pyr synapses to 40 ms. In the low ChC percentage condition, this further reduced the 40/40 amplitude while increasing the 20/20 measure. Additionally, a 20/40 peak emerged. However, there was also a strong reduction of the 30/30 measure (Figure 7). Overall, these results here are very similar to the results from the simulations where only IPSC times at BC-Pyr synapses were modified, suggesting a minor role of ChCs in the generation of ASSR deficits in schizophrenia.

**Figure 7:**
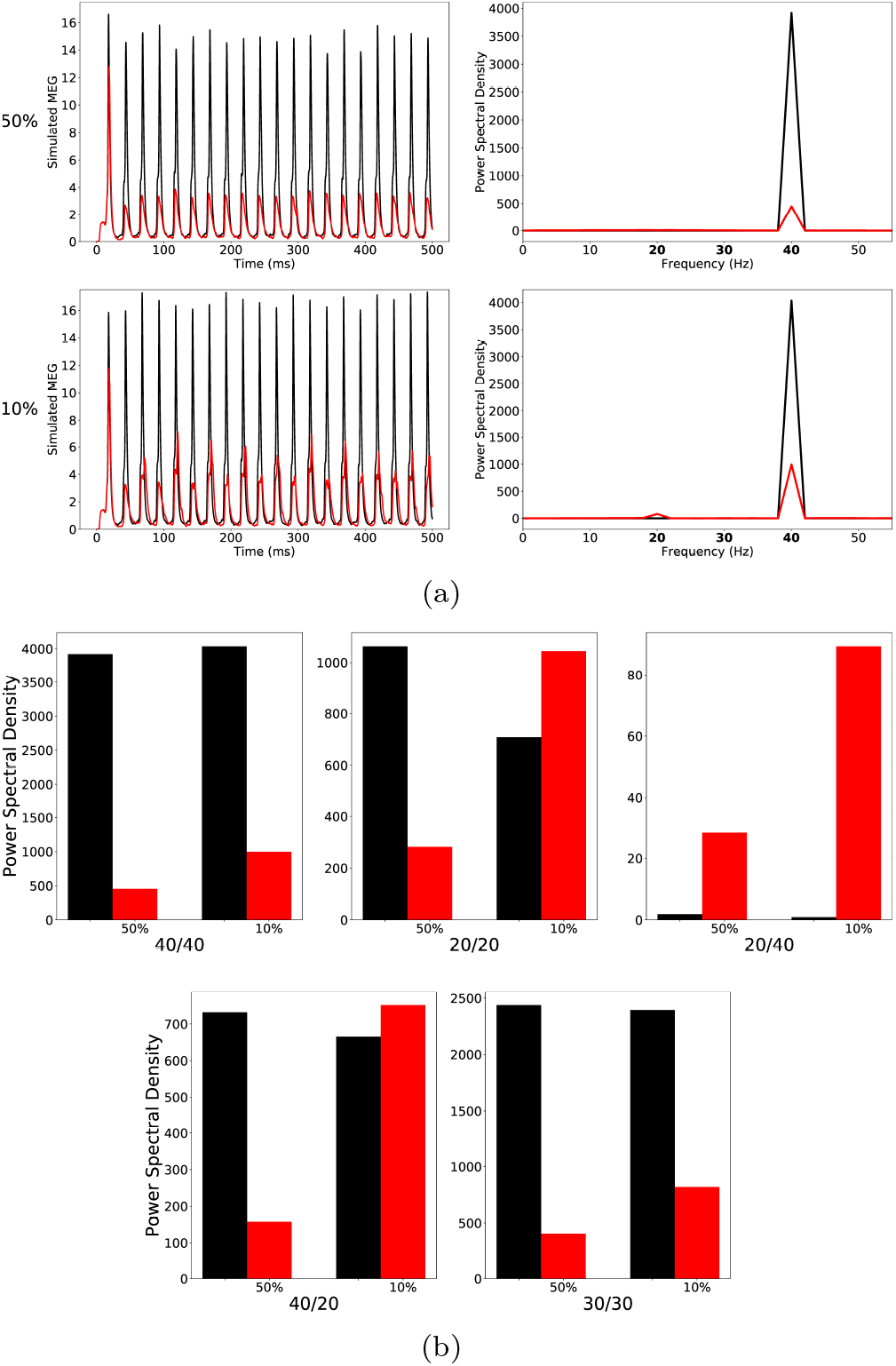
(a) left row: Raw simulated MEG signals for high (50%) and low (10%) percentage of chandelier cells among total inhibitory cells for a drive frequency of 40 Hz and the two network conditions (‘control’: black traces; ‘schizophrenia-like’: red traces). Right row: Power spectral density of the MEG signals of the left row. (b) Comparison of the 5 entrainment measures between control (black) and schizophrenia-like (red) for high and low ChC percentages. Note that ‘schizophrenia-like’ here means that IPSC decay times at **chandelier** cell synapses were increased from 8 ms to 28 ms and at **basket** cell synapses were increased from 8 ms to 20 ms. All traces here depict the mean signal averaged over 20 trials with different realisations of the Poisson process underlying the generation of noise. Importantly, averaging was performed in the time domain prior to a transformation into the frequency domain via Fourier transform, which is also the standard procedure in experiments.

As Figure 6, but the IPSC decay times at **chandelier** cell synapses were increased to 40 ms and at **basket** cell synapses to 28 ms to model ‘schizophrenia-like’ networks. (a) left row: Raw simulated MEG signals for high (50%) and low (10%) percentage of chandelier cells among total inhibitory cells for a drive frequency of 40 Hz and the two network conditions (‘control’: black traces; ‘schizophrenia-like’: red traces). Right row: Power spectral density of the MEG signals of the left row. (b) Comparison of the 5 entrainment measures between control (black) and schizophrenia-like (red) for high and low ChC percentages. All traces here depict the mean signal averaged over 20 trials with different realisations of the Poisson process underlying the generation of noise. Importantly, averaging was performed in the time domain prior to a transformation into the frequency domain via Fourier transform, which is also the standard procedure in experiments.

## Discussion

Here, we explored the contribution of slower IPSC dynamics at ChC-Pyr synapses to gamma and beta range ASSR deficits in patients with schizophrenia using a model cortical network. Generally, at realistic, low ratios of ChCs to BCs, increased IPSC decay times at ChC synapses alone hardly had any influence on ASSRs. At low ChC/BC ratios, increased decay times at BC-Pyr synapses, on the other hand, strongly altered the spectral composition of ASSRs, replicating several characteristics found experimentally. For a moderate increase (i.e. *τ_BC_* =20ms), we found a strong decrease of the 40/40 measure, an increase of the 20/20 measure and an increase of the 20/40 measure, although it remained low (Figure 3). This suggests that this change shifts the preferred frequency of the network from the gamma range more towards the beta range and gives an explanation of the reduced gamma seen across many experiments. Furthermore, the 30/30 measure remained virtually unaffected. However, it did not explain the strong 20/40 component seen by Vierling-Claassen et al. [41]. For a stronger increase (i.e. *τ_BC_*=28 ms), we found an even stronger decrease of the 40/40 measure, a further increase of the 20/20 measure and a much stronger increase of the 20/40 measure (see Figure 4). However, we also saw an increase of the 40/20 measure, contradictory to the gamma to this beta shift hypothesis. Furthermore, the 30/30 measure becomes greatly reduced (Figure 4 (b)), which is not seen experimentally. Looking at the power spectra for 40 Hz and 30 Hz drive for increasing *τ_BC_*, we see that with longer decay time the 20/40 measure gets stronger but at the same time the change to the network increasingly effects the 30/30 measure (Figure 5). Lastly, when looking at the effect of increased decay times of ChC-Pyr synapses in a network where also BC-Pyr synapse dynamics are slowly decaying, we see that the additional contribution of ChC-mediated effects is very weak for different combinations of decay times *τ_BC_* and *τ_ChC_* (see Figures 6 and 7). One interesting effect, however, was the smaller 20/40 measure here compared to the condition where only BC-Pyr synapses had longer decay times (compare Figures 4 and 7).

Overall, our findings suggest that at realistic, low ratios of ChCs to BCs in a microcircuit, gamma and beta range changes to the spectral composition of ASSRs as seen in SCZ patients, can be well explained by an increase of GABAergic decay times at BC-Pyr synapses. While postmortem studies indicate that the reduction in GAT1 in SCZ patients preferentially affects ChCs [21], this might simply reflect easier detection due to the specifc morphology of ChC synaptic cartridges compared to BC synapses and not a general difference in GAT1 reduction between the two subtypes. Nevertheless, prolonged IPSC decay times as a result of reduced GAT1 levels requires a high synapse density [29] and, thus, it is not immediately clear if a GAT1 reduction would be sufficient to cause a prolonged IPSC time course at BC synapses. Interestingly, however, repeated activation together with a reduction in GAT1 also resulted in an increase in IPSC decay times in the study of [29]. Therefore, increased IPSC times might also be present at BC synapses, especially in a gamma and beta entrainment setting where the network is driven with a relatively high frequency.

Our results demonstrate that prolonged IPSC decay times at BC-Pyr synapses robustly lead to a weakening of gamma synchronization, especially during gamma drive but also during beta drive. This is consistent with a large body of experimental evidence reporting reduced gamma band power during ASSR paradigms [19, 41, 18, 36, 39, 22, 4, 34, 32, 23, 37] but also during various sensory (e.g. a visual Gestalt task [33]), and cognitive tasks (e.g. a working memory task [5]). We also found an increase of beta synchonization during beta drive, and, although in most conditions still very small, increased beta synchronization during gamma drive. This reproduces the findings of Vierling-Claassen et al. [41] that slow dynamics at FS PV^+^ interneurons might not only desynchronize gamma oscillations but also shift the preferred frequency of cortical networks from gamma to the beta band. However, our results further demonstrate that, most likely, BCs are the main interneuron subtype causing this shift and not ChCs, as hypothesized by Vierling-Claassen et al. [41]. Furthermore, we found that while longer decay times at BC-Pyr synapses led to a stronger and more robust peak of the 20/40 measure, they also introduced a stronger reduction of the 30/30 measure. Given that studies have consistently reported that the 30/30 measure remains unaffected in SCZ patients [19, 41, 37], together with the fact that the strong 20/40 component has not been reported by experimental studies except for Vierling-Claassen et al. [41], our results suggest that a strong 20/40 component might not be a core ASSR deficit in patients with schizophrenia. This is further underpinned by the finding that alterations at ChCs together with BC alterations, led to a reduction of the 20/40 measure compared to the condition where only BC-Pyr had prolonged decay times.

While the results presented here suggest a modulatory influence of ChC alterations on gamma and beta oscillations at most, we have to note that the employed model has substantial limitations. Our model, for example, does not explore the possibility that, due to the location directly at the axon initial segment, ChCs might exert significantly stronger control over PC firing, which in turn might counterbalance their small numbers. Although it is not intuitively clear how a combination of prolonged IPSC times and a potentially depolarizing effect of ChCs [44, 45, 46] result in entrainment deficits comparable to experimental findings, this has not yet been explored. Furthermore, our model does not include NMDA receptors, which have been linked to gamma range deficits in schizophrenia [17, 14]. We therefore haven’t explored the potential consequences of NMDA hypofunction differently affecting BCs compared to ChCs within this model of prolonged IPSC times. Finally, the theta neuron model is a simple and abstract neuron model and does not capture electrophysiological differences between BCs and ChCs found experimentally [30]. Interestingly, genetic variants of risk genes for schizophrenia include a variety of genes coding for ion channels or ion transporters, which have been shown to alter cell excitability [24, 26] and network oscillations [25]. Such variants might potentially amplify electrophysiological differences between BCs and ChCs and thereby change their relative roles in the generation of gamma/beta oscillations.

In summary, our findings suggest that ChCs are likely not strongly involved in gamma/beta range ASSR deficits in schizophrenia, and, therefore future modelling efforts should focus on alterations at BC-Pyr synapses. Furthermore, ChC-Pyr synapses might not represent a valuable target for therapeutic interventions aiming at restoring the gamma ASSR in patients with schizophrenia.

